# The estrogen receptor α cistrome in human endometrium and epithelial organoids

**DOI:** 10.1101/2022.04.19.488787

**Authors:** Sylvia C Hewitt, San-pin Wu, Tianyuan Wang, Mita Ray, Marja Brolinson, Steven L. Young, Thomas E. Spencer, Alan DeCherney, Francesco J DeMayo

**Affiliations:** Pregnancy & Female Reproduction, RDBL, NIEHS, Research Triangle Park, NC 27709; Integrative Bioinformatics Support Group, NIEHS, Research Triangle Park, NC 27709; Program in Reproductive and Adult Endocrinology, NICHD, Bethesda, MD 20847; Department of Obstetrics and Gynecology, University of North Carolina at Chapel Hill, Chapel Hill, North Carolina 27599; Department of Obstetrics, Gynecology and Women’s Health, University of Missouri, Columbia, Missouri 65211

## Abstract

**Context:** Endometrial health is impacted by molecular processes that underlie estrogen responses.

**Objective:** To define estrogen regulation of endometrial function by integrating the estrogen receptor alpha (ESR1) cistrome and transcriptome of endometrial biopsies taken from the proliferative and midsecretory phases of the menstrual cycle and hormonally stimulated endometrial epithelial organoids.

**Design:** ESR1 ChIPseq and RNAseq were performed on proliferative or mid-secretory endometrial biopsies and on hormone treated organoid cultures.

**Setting:** Endometrial samples were obtained from volunteers at outpatient research clinics for ChIPseq and for organoid culture.

**Patients or Other Participants:** Participants were fertile, reproductive aged women with normal cycle length, and without any history of infertility or irregular cycles. In total, 5 new endometrial biopsies obtained from 5 women were used in this study and were analyzed together with previously published cycle stage endometrial RNAseq data.

**Intervention(s):** There were no interventions in this study.

**Main Outcome Measure(s):** The cycle stage specific ESR1 binding sites and gene expression identification of human endometrium and organoid cultures were integrated with changes in gene expression.

**Results:** Genes with ESR1 binding in whole endometrium were enriched for chromatin modification and regulation of cell proliferation. The distribution of ESR1 binding sites in organoids was more distal to the gene promoter when compared to primary endometrium. Organoid estrogen/ESR1 candidate target genes impacted formation of cellular protrusions, and chromatin modification,

**Conclusions:** Analysis of the ESR1 cistromes and transcriptomes from endometrium and organoids provides important resources for understanding how estrogen impacts endometrial health and function.

## Introduction

Uterine function depends on the precise coordination of ovarian steroid hormone signaling. The actions of estrogen and progesterone, via their cognate receptors, the estrogen receptor ESR1 and progesterone receptor, PGR, regulate the ability of the uterus to support embryo implantation and fetal growth. Alterations in estrogen action can result in diseases of the uterus such as endometriosis and endometrial cancer. Understanding how these hormones regulate uterine biology is critical to understanding their roles in uterine function and dysfunction.

The uterus contains multiple types of cells arranged into layered compartments with an interior lumen that is lined by epithelial cells (1). Glandular structures with specialized epithelial cells emanate from the lumen into the uterine stromal tissue and secrete cytokines to facilitate establishment and maintenance of pregnancy. Underlying the single-cell epithelial lumen and glands are stromal fibroblasts, blood vessel endothelium and bone-marrow derived immune cells. The stromal fibroblasts undergo a marked structural and functional transformation, beginning before implantation in the mid-secretory phase, into decidual cells, in a process termed decidualization. The endometrium includes the stromal and epithelial compartments and when pregnancy does not occur, the corpus luteum fails and ovarian hormones decrease, causing the endometrium to be shed by menstruation (2). The hormone dependent endometrial stages are classified based on histological examination as the proliferative (pre ovulatory, estrogen dominant) and secretory (post ovulatory, progesterone dominant) phases. The mid-secretory phase is the time when the endometrium is receptive to embryo implantation (3).

During the proliferative phase of the menstrual cycle, estrogen induces uterine cell growth and endometrial tissue thickening, and acts to increase the progesterone sensitivity of the endometrial cells (3). Situations or conditions that perturb estrogen signaling are detrimental to normal uterine development and function. A dramatic example was provided when prescription of the potent synthetic estrogen diethylstilbestrol (DES) to pregnant women resulted in profound in utero developmental alterations in uterine structures of female fetuses (4). “DES daughters” have an increased risk for infertility, ectopic pregnancy, miscarriage, and clear cell adenocarcinoma (5,6). Alternatively, women who have mutations in their estrogen receptor causing estrogen insensitivity or who are unable to produce estrogen due to mutations in the enzyme aromatase have an abnormally small uterus with a thin endometrial lining and are infertile. (7-11). Chronic exposure to estrogen without the growth-inhibiting effects of progesterone is a major contributing factor to endometrial hyperplasia and subsequent cancer (12,13). Estrogen also facilitates the progression of endometriosis by increasing growth of lesions and can impact the severity of endometriosis symptoms (14). Finally, we reported previously that thin endometrium, associated with poor fertility treatment outcomes, is associated with defects in estrogen signaling (15). Therefore, understanding mechanisms for optimal estrogen signaling is a key element in maximizing women’s health.

Estrogen impacts endometrial cells via nuclear estrogen receptors (ESR1 and ESR2) (16,17), that act as estrogen dependent transcription factors by binding to estrogen and to specific DNA motifs (GGTCAnnnTGACC), called estrogen responsive enhancers (EREs). ESR binding to EREs in accessible regions of DNA and interaction with chromatin modifiers leads to alteration in transcriptional rates of estrogen responsive genes (16). Utilizing chromatin immunoprecipitation and next generation sequencing (ChIPseq), the locations of all ESR1-chromatin interaction within cells or tissues can be ascertained, defining the ESR1 cistrome (18,19). These analyses indicate that receptor interactions are frequently in enhancer regions that are far (in linear sequence length) from the transcriptional start of the responding genes, highlighting the importance of the 3-dimensional arrangement of the chromatin structure by which the ESR1/ERE is brought into contact with its target genes.

Much of the molecular detail of estrogen action has been defined using *in vitro* cell line models or *in vivo* in experimental animal models. Although there are many similarities between mechanisms in women and these model systems, application to women’s health is best done by determining the key details in women. *In vivo* and *ex vivo* studies in women are, of course, limited by ethical and practical considerations, and so studies done in women need to be focused on the most important aspects of endometrial function.

Endometrial epithelial cells provide the first barrier to successful embryo implantation (21), and thus optimal function of endometrial epithelium is key to reproduction. However, studies focused on details intrinsic to the epithelial cells are particularly challenging for two main reasons: 1) Biochemical characterization of clinical samples is limited due to the invasive process of endometrial biopsy required to obtain samples from healthy volunteers. 2) Although uterine stromal cells can be cultured and studied, epithelial cells have proven difficult to culture in a model that recapitulates their biological environment and function. The recent development of organoid culture has provided a system in which endometrial epithelial cells can be grown and studied *in vitro* (21). Previous work has characterized ESR1 interactions with chromatin in cultured breast cancer (22) and endometrial cancer cells (23), endometrial tumors (24) and endometrial biopsies from infertile women treated with clomiphene (15). Here, we describe the profile of estrogen receptor interaction with chromatin in human endometrial samples from healthy volunteers at the estrogen dominant proliferative phase or the progesterone dominant mid-secretory phase. Further, we cultured epithelial cells from endometrial biopsies in organoids to describe details of estrogen response in an *in vitro* system. The transcriptional profiles and ESR1 binding sites derived from the isolated epithelial cells are revealed, and then are compared to observations derived from whole endometrium, increasing our understanding of the estrogen receptor mediated processes that are intrinsic to endometrial epithelial cells. Together, our study allows examination of ESR1 interactions with endogenous enhancers and promoters that mediate transcriptional responses within normal endometrial tissue.

## Materials and Methods

### Ethics statement

This project was executed in accordance with the federal regulation governing human subject research. The study was reviewed and approved by the Committee for the Protection of Human Subjects at the University of North Carolina at Chapel Hill IRB under file #05-1757 or NIH Institutional Review Board 99CH0103. Informed consent was obtained from all patients before their participation in this study.

### Human endometrial samples

Healthy women, ages 19 to 34 years, with a regular inter-menstrual interval between 25 and 35 days and no history of infertility or pelvic disease were invited to participate. Exclusion criteria were the following: a) an inter-menstrual interval that varied by more than three days, b) use of medication known to affect reproductive hormones or fertility within 60 days prior to enrollment, c) chronic disease, d) a body mass index > 29.9 or < 18.5, and e) history of infertility, defined as a failure to conceive for one year or longer despite regular intercourse without contraception.

All subjects underwent an endometrial biopsy, taken from the mid-fundus, in the office, using a Pipelle™ suction curette. For isolation of epithelial cells for organoid culture, endometrial biopsies were taken from volunteers and processed immediately. For ESR1 ChIPseq, biopsies were obtained from proliferative or mid-secretory phases and frozen.

#### ESR1 ChIPseq

Frozen endometrial samples or fixed, frozen organoid pellets were shipped to Active Motif Inc (Carlsbad CA) for Factor Path ESR1 ChIP (ERα antibody 06935, EMD Millipore, Temecula, CA) and library preparation. Libraries were shipped to NIEHS and sequenced in the NIEHS Genomics Core Laboratory. Raw ChIP-seq reads were processed and aligned to the human reference genome hg38 using Bowtie (25). The reads were de-duplicated, and peaks were called using MACS2 (26). The mergePeaks function of HOMER (27) was used to find shared ESR1 peaks in donor1 and donor2 organoid samples and to make a Venn diagram of ESR1 peaks in proliferative vs mid-secretory endometrium. Peak Annotation and Visualization (PAVIS; (28)) was used to compare the locations of ESR1 peaks relative to genes. Known motifs in ESR1 peaks were identified using HOMER findMotifs. Heatmap plots of ESR1 signal centered on organoid ESR1 peaks were created using EaSeq (29). EaSeq was also used to locate the nearest gene TSS and TES to each ESR1 peak as a way to identify peaks within 100 kb of genes. Data is deposited in GEO GSE200807.

### Endometrial RNAseq analysis

RNAseq FPKM values from our previous study (30) for proliferative or mid-secretory whole endometrium or isolated epithelium RNA were used (GSE132713). Whole endometrium was analyzed using ANOVA and List Manager in Partek Genomics Suite software (Partek Inc, St Louis MO) to find differentially expressed genes (DEG, 2-fold, FDR p-value<0.05) and filter the DEG list to include DEG within 100kb of ESR1 peaks. This filtered gene list was imported into Ingenuity Pathway Analysis (IPA, Qiagen, Redwood City, CA) for Core Analysis.

### Organoid Culture and Treatment

Organoids were derived from fresh donor endometrial biopsies collected and washed in wash medium (DMEM/F12 (Gibco, Amarillo, TX) + 1x Anti-Anti (Gibco)), minced, and digested in 20 ml 0.4 mg/ml Collagenase V (Sigma)+2.5 mg/ml Dispase II (Sigma) dissolved in wash medium at 37 C for 50 minutes. Digestion was stopped by adding 10% fetal bovine serum (FBS) diluted in wash medium. Remaining tissue debris was allowed to settle, and the suspended tissue digest was passed through a 100 µm Cell Strainer (Falcon, Brookings SD) and washed with wash media. Epithelial cells were backwashed from the cell strainer, centrifuged at 232 RCF for 10 minutes, resuspended and washed. The final pellet was resuspended in Matrigel (Corning, Corning NY) to make the final concentration 90% Matrigel, 3-4 25 µl drops were plated in each well of 12-well, plate and overlayed with 700 µl of expansion media (Advanced DMEM/F12 (Gibco) containing B27 (minus vitamin A, Gibco), Insulin-Transferrin-Selenium (ITS, Gibco), Primocin (Invitrogen), Glutamax (Gibco), N2 supplement (Gibco),1.25 mM N-acetyl-L-cysteine (Sigma), 500 nM A83-01 (Tocris, Bristol, UK), 50 µg/ml human HGF (Peprotech, Cranbury, NJ), 500 µg/ml human EGF (Peprotech), 100 µg/ml human FGF-10 (Peprotech), 500 µg/ml human Rspondin-1 (Peprotech), 100 µg/ml human Noggin (Peprotech), 10 nM Nicotinamide (Sigma). Media was replaced every 2-3 days. Once organoids formed and grew to fill the Matrigel drop (see Fig. S2b), they were passaged by resuspension in Advanced DMEM/F12+B27, ITS, Primocin, and Glutamax, centrifuged and washed to release organoids form Matrigel and either frozen in 10% DMSO (Sigma) in FBS and stored in liquid nitrogen, or resuspended and plated in fresh Matrigel. For hormone treatments (see graphic in Fig S2a), 4 days after organoids were plated, media was replaced with expansion media that had N2 omitted and contained either 0.1% ethanol (V) or 10 nM estradiol (E2, Steraloids, Newport, RI). For RNA isolation, 2 (d6), 5 (d9) or 8 (d12) days later fresh media containing V or E2 was added, and organoids were isolated 6 hours later. For the ESR antagonist treatment (Fig. 2b), on day 8, and again on d9 fresh media containing V, E2, or 10 nM E2+1 µM ICI, 780 (Tocris) was added, and organoids were collected 6 hours after the media change on d9. For ChIPseq, on d9, fresh media containing 10 nM E2 and 1 µM medroxyprogesterone acetate (Sigma) was added, and organoids were isolated one hour later. The progesterone was included to for planned future PGR ChIPseq analysis of remaining chromatin.

### RNA Isolation and Analysis

RNA was isolated from organoids using Trizol Reagent (Invitrogen) according to the manufacturer’s instructions. For RT-PCR, cDNA was prepared using Superscript II (Invitrogen) with Random Hexamers (Invitrogen), as previously described (31,32). Sequences of primers (Sigma) used are: ESR1 F-CTGCAGGGAGAGGAGTTTGTGT R-TCCAGAGACTTCAGGGTGCT, IHH F-GACCGCGACCGCAATAAGTA R-TGGGCCTTTGACTCGTAATAC, GAPDH F-ATGGGGAAGGTGAAGGTCG R-GGGGTCATTGATGGCAACAATA, GREB1 F-ATGGGAAATTCTTTACGCTGGAC R-CACTCGGCTACCACCTTCT, PGR F-GACGTGGAGGGCGCATAT R-AGCAGTCCGCTGTCCTTTTCT. PCR was done using SsoAdvanced Universal SYBR green Supermix (BioRad) with a CFX instrument (Biorad).

For RNAseq, RNA (3 replicates each of V or E2 treated samples from donor 1 and donor 2) was DNAse treated and cleaned up using the RNeasy Mini kit (Qiagen) or the RNA Clean and Concentrator 5 kit (Zymo, Irvine CA). RNA was submitted to the NIEHS sequencing core for library preparation and paired end sequencing using Illumina’s Ribo-Zero Gold kit for donor 1 RNA and Illumina Stranded mRNA Prep for donor 2. Raw data was filtered to remove low quality reads, mapped to hg38 using Tophat (33) and de-duplicated using Picard tools (2.18.15; “Picard Toolkit.” 2019. Broad Institute, GitHub Repository. https://broadinstitute.github.io/picard/; Broad Institute). BAM files were imported into Partek Genomics Suite for analysis. RPKM for RefSeq genes were determined, and the maximum RPKM value from any sample for each gene (MAX) was listed. We noted differences between donor 1 and donor 2 datasets that reflect differences in RNA and library quality. For this reason, we analyzed the datasets from each donor separately, and then compared the DEG, so that each would be relative to its own V control. Genes for which the MAX< 1% of the mean RPKM for the whole dataset were filtered out (<0.02 for donor 1, <0.002 for donor 2). Values of 0 for V samples were replaced with a value of 0.05% of MAX to allow calculation of E2/V. After PCA, it was noted that donor 1 V-1 and donor 1 E2-1 samples were outliers, potentially due to low reads, and were excluded from further analysis. Partek ANOVA was used to determine E2 vs V fold changes, and the List Manager function was used to find DEG (2-fold, FDR p-value<0.05). Donor 1 or donor 2 DEG data were analyzed individually using IPA, and analyses were compared using the Comparison function. Partek Venn diagrams were used to construct a list of all DEG in either donor to combine E2 vs. V fold changes of the datasets for visualization. Data is deposited in GEO (GSE200807).

## Results

### Endometrial ESR1 Cistrome

We previously described PGR interaction with chromatin (ChIPseq) together with transcriptomic (RNAseq) analysis in proliferative or mid-secretory stage endometrial biopsies from healthy volunteers (30). Here, we have extended those findings by evaluating ESR1 binding using the same techniques in similar samples. In the whole endometrial samples, there are 4-fold more ESR1 peaks in the proliferative sample (35,156 peaks) than the mid secretory sample (8,688 peaks), consistent with the estrogen dominant proliferative phase (Fig. 1a). We examined the locations of the ESR1 peaks relative to annotated genes. ESR1 peaks were distributed comparably in the proliferative and mid-secretory endometrium (Fig. 1b), with about half of peaks located at genes (49% of proliferative and 52% of mid secretory ESR1 peaks at 5’UTR, exons, introns and 3’ UTR). Motif analysis of the ESR1 peaks indicates that ERE and HOX motifs are enriched in both proliferative and mid-secretory samples (Fig. 1c). ERE is the most significantly enriched motif in proliferative samples (Fig. 1c), and HOX motifs are the top enriched motifs in mid-secretory ESR1 peaks (Fig. 1c). In addition, bZIP/AP1 and bHLH motifs were seen in both proliferative and mid-secretory ESR1 binding peaks.

**Figure 1.**
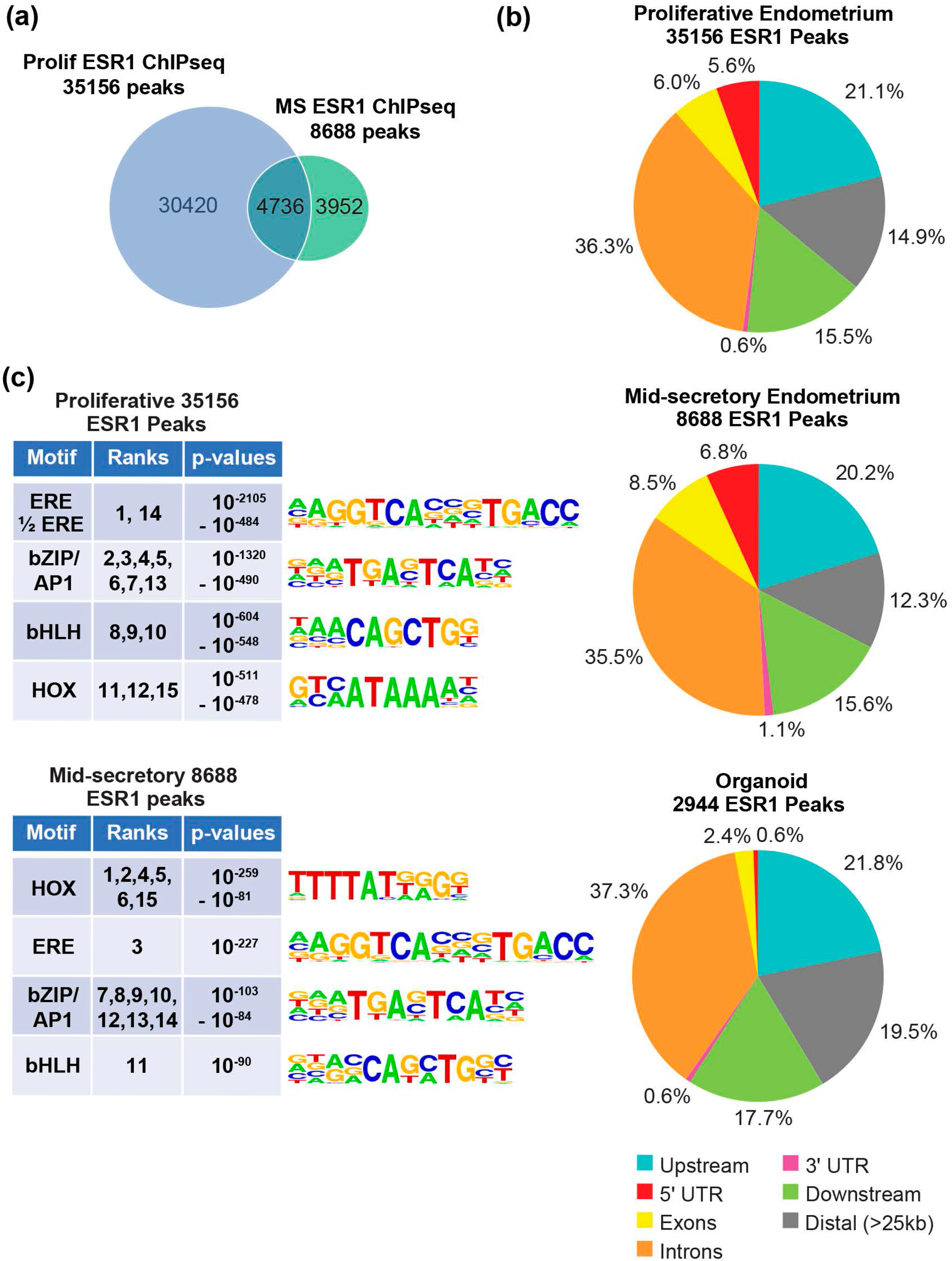
ESR1 ChIPseq of proliferative and mid-secretory endometrial biopsies. a. Venn diagram comparing number of ESR1 peaks in proliferative (Prolif) or mid-secretory (MS) endometrial biopsies. b. Locations of ESR1 peaks from proliferative or mid-secretory endometrium or from organoids, relative to annotated genes (RefSeq). c. Summaries of the top 15 HOMER known motifs enriched in ESR1 peaks from proliferative or mid-secretory endometrial biopsies. The ranks of each motif, as determined by p-value, along with the range of p-values covered by the ranks are indicated. Motif logos are shown.

To further evaluate the impact of ESR1 chromatin interactions on uterine functions, we determined the ESR1 binding sites that are within <100 kb of an annotated Refseq gene in the proliferative (31,814 peaks) or mid-secretory (7,931 peaks) samples (Fig. S1). Next, we analyzed the endometrial transcriptome, which we previously described (30), by identifying genes that were expressed in proliferative and mid-secretory RNA (FPKM≥1 in at least one sample) and determining those that were differentially expressed between proliferative and mid-secretory phases (1628 DEG 2-fold, FDR p-value<0.05; Table S1a). Genes that differ between the estrogen dominant proliferative and progesterone dominant mid-secretory phases include ESR1 targets that are either increased or decreased by estrogen. We then filtered the DEG to include genes that are <100 kb from the proliferative (902 genes; Table S1b) or mid-secretory (394 genes) ESR1 peaks, respectively (Fig. S1a) as genes that are candidate ESR1 downstream targets.

We focused on 902 proliferative endometrium DEG that are candidate ESR1/estrogen targets and used pathway analysis (Ingenuity Pathway Analysis) to assess their impact on biological functions and signaling (Table 1 and Table S1c,d,e). Our analysis indicates impacts on multiple overlapping processes including estrogen and progesterone signaling, proliferation, chromatin modification, transcription, endometrial physiology associated signals, lipids, growth factor/receptor tyrosine kinase signaling, signaling via other receptors, and hematological system development.

**Table 1:**
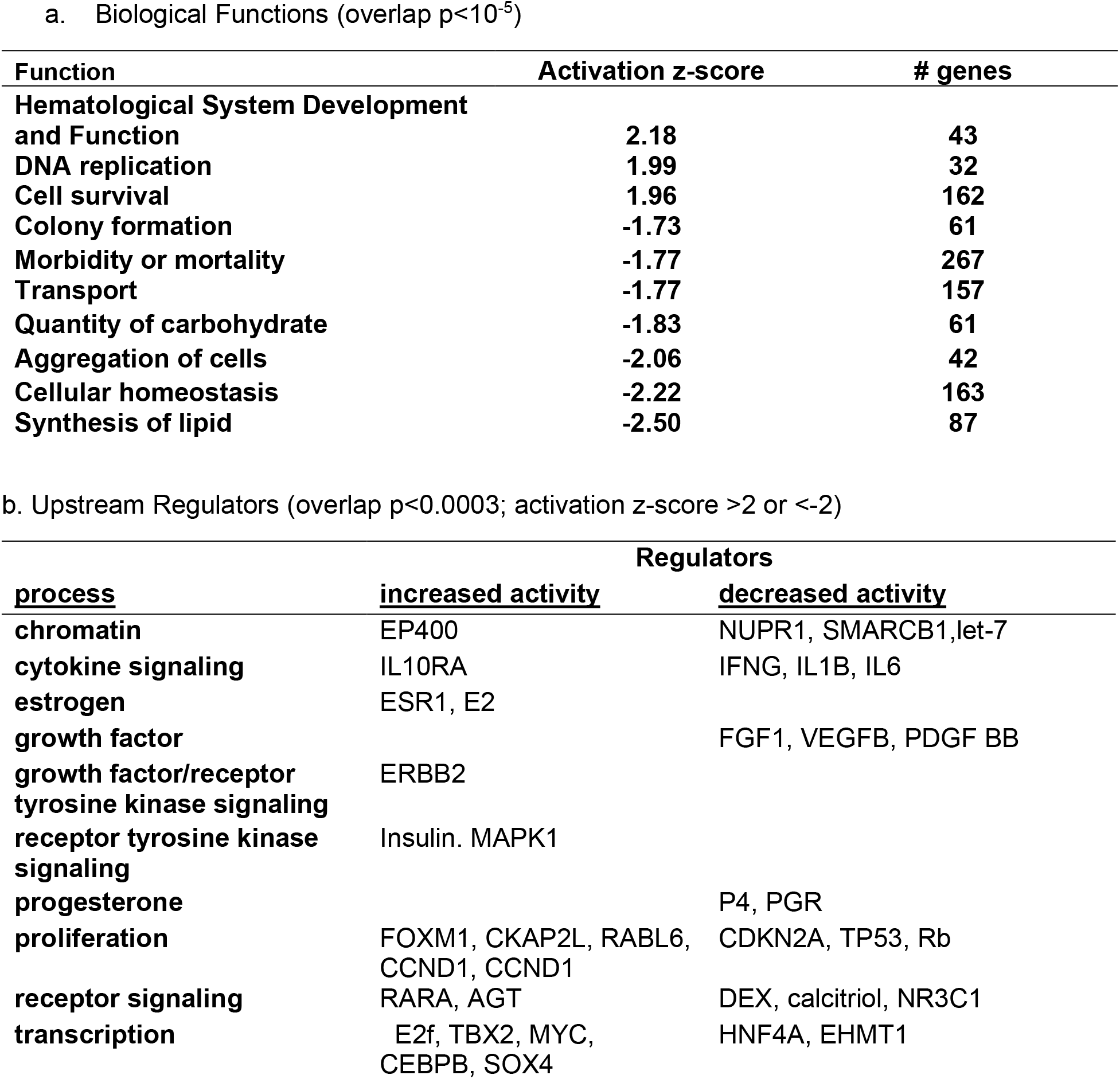
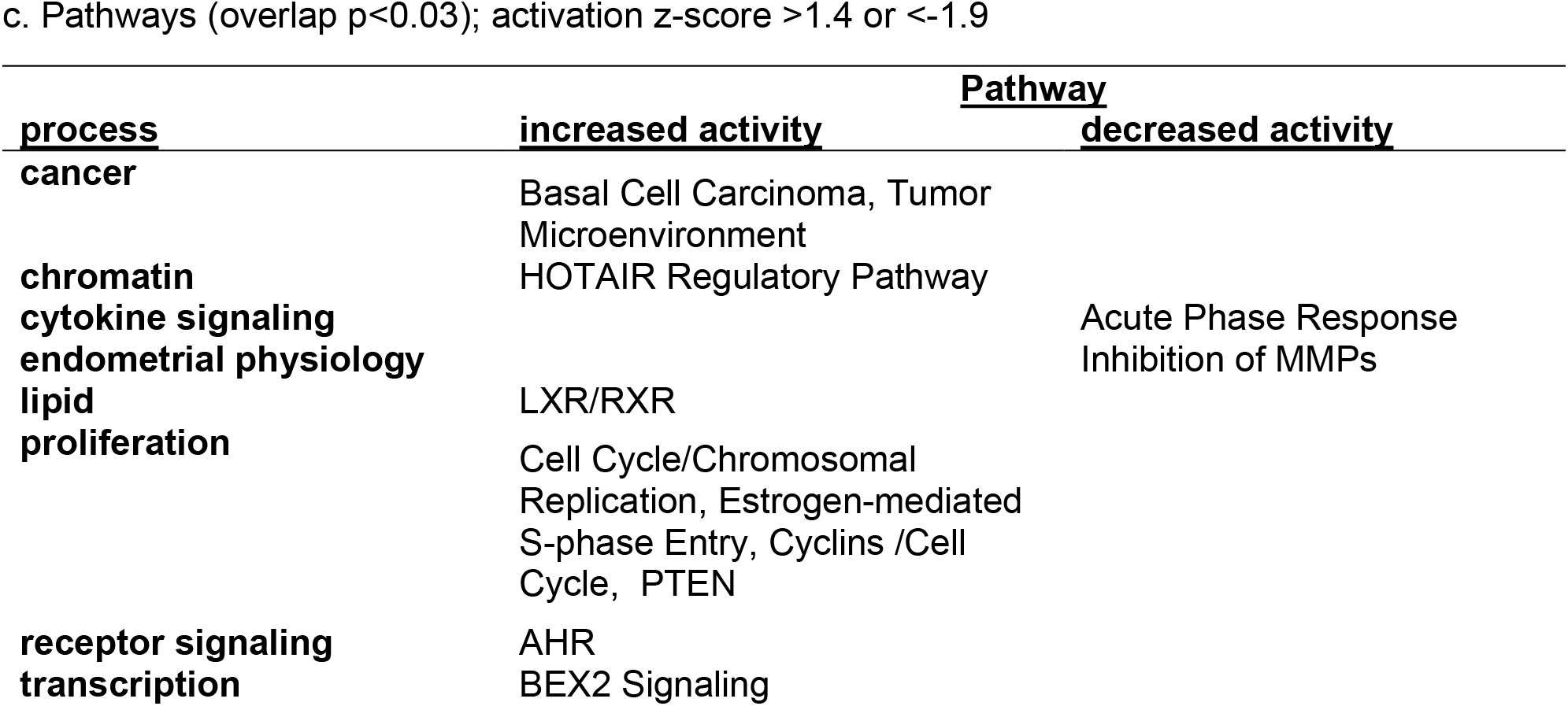
Functions (a), Upstream Regulators (b) and Pathways (c) Enriched in 902 Proliferative vs Mid-secretory Differentially Expressed Genes That are Less than 100 kb From a Proliferative ESR1 ChIPseq Peak

We noted increased estrogen/ESR1 activity and decreased progesterone/PGR activity (Table 1b), as expected of genes that differ between estrogen dominant proliferative phase vs. the progesterone dominant mid-secretory phase and validating that our approach identifies potential ESR1 regulated endometrial pathways.

Proliferation associated functions, signaling pathways and regulators are enriched, including DNA replication (Table 1a) chromosome replication, E2-mediated S phase entry, cyclins and cell cycle (Table 1c) together with the regulators mitotic spindle component Cytoskeleton associated protein 2 like (CKAP2L), cell growth factor RAB, member RAS oncogene family like 6 (RABL6), cyclin CCND1, mitosis factor Forkhead box M1 (FOXM1), and inhibition of cell cycle inhibitors Rb tumor suppressor and Tumor protein 53 (TP53) and TP53 stabilizer Cyclin dependent kinase inhibitor 2A (CDKN2A; Table 1b). Altogether, these indicate increased proliferation, which corresponds to estrogen stimulated endometrial growth that is characteristic of the proliferative phase.

The gene set impacts chromatin modifiers including activation of HOX transcript antisense RNA (HOTAIR; Table 1c), a long non-coding RNA associated with multiple cancers that acts as a guide and scaffold to deliver Polycomb repressive complex 2 (PRC2) chromatin modifiers to specific gene loci (34), histone acetyl transferase (HAT) complex factor E1A binding protein p400 (EP400), and inhibition of micro RNA let-7, which targets HOTAIR. Additional chromatin regulators affected include SWI/SNF component SWI/SNF related, matrix associated, actin dependent regulator of chromatin, subfamily b, member 1 (SMARCB1), which removes repressive chromatin marks, and HAT activity factor Nuclear protein 1, transcriptional regulator (NUPR1; Table 1b).

Candidate Estrogen/ESR1 gene responses were consistent with impacts on activities of multiple transcription factors (increased activity of transcription factor brain expressed X-linked 2 (BEX2) pathway (Table 1c), increased activity of E2f, T-box transcription factor 2 (TBX2), MYC proto-oncogene, CCAAT enhancer binding protein beta (CEBPB), and SRY-box transcription factor 4 (SOX4), inhibition of transcription factor hepatocyte nuclear factor 4 alpha (HNF4A) and transcription inhibitor Euchromatic histone lysine methyltransferase 1 (EHMT1; Table 1b)). Effects on chromatin and transcription align with estrogen/ESR1 having roles in chromatin modifications that influence accessibility of transcriptional mediators.

Several signaling pathways known to be important in endometrial physiology and pathophysiology were enriched. For example, activity of the AKT inhibitor and tumor suppressor, phosphatase and tensin homolog (PTEN) was increased. PTEN has an essential role as a cell cycle checkpoint regulator to prevent premature mitosis and resulting gene mutations (35) and is frequently inactivated in endometrial tumors (36). Further, estrogen inhibition of endometrial cytokine signals are reflected by decreased acute phase response (Table 1c) and cytokine signaling (Interleukin 1B (IL-1B), IL-6, Interferon γ (IFNG), Table 1c) and increased Interleukin 10 receptor subunit A (IL-10RA) signaling (Table 1c) in the proliferative phase ESR1-associated genes, aligning with suppression of cytokine signaling during the proliferative phase and later activation associated with implantation (37,38). There are indications that ESR1 signaling affects lipids, as reflected by decreased activity of lipid synthesis (Table 1a) and Liver X receptor/Retinoid X receptor (LXR/RXR) activation (Table 1c).

Several mediators of growth factor/receptor tyrosine kinase (RTK) pathways are impacted as well. The RTK Erb-b2 receptor tyrosine kinase 2 (ERBB2) is activated, as is the downstream modulator Mitogen-activated protein kinase 1 (MAPK1; Table 1b). The activity of growth factors platelet derived growth factor BB (PDGFBB), vascular endothelial growth factor B (VEGFB) and Fibroblast growth factor 1 (FGF1) are decreased (Table 1b). Signaling via other receptors is also evident, including activation of Aryl hydrocarbon receptor (AHR; Table 1c) and Retinoic acid receptor a (RARA; Table 1b), and inhibition of calcitriol (vitamin D agonist) and dexamethasone (glucocorticoid receptor agonist; Tables 1b and c).

Hematological system development and function was enriched (Table 1a), which includes enhanced signaling from angiotensin (AGT; Table 1b). This observation aligns with studies indicating angiogenesis, vascular remodeling and regulation of blood flow in the endometrium and placenta are essential processes regulated by the renin-angiotensin system (39-43). Overall, our analysis of ESR1 associated candidate estrogen regulated processes aligns with estrogen’s multiple roles in the proliferative phase endometrium, both previously described and novel.

### Estrogen Responses of Organoids

Development of organoid models derived from epithelial cells isolated from endometrial biopsies has provided an *in vitro* system in which to manipulate and study hormone signaling mechanisms (44-46). E2 responses were evaluated in organoids cultured as previously described by Fitzgerald et al (44) with some modification. Organoids were plated and allowed to form for 4 days, then treated with E2 or V for 2, 5, or 8 more days (day 6, day 9, day 12; Figure S2a). Samples were collected 6 hours following a change to fresh media containing V or E2 (see methods section for detail) to capture acute responses. Organoids derived from endometrial samples obtained from three different donors were compared to assess individual donor variability. The levels of transcript for three estrogen regulated endometrial genes (*PGR, IHH*, and *GREB1*) were measured. All three genes were robustly induced by E2 in all three donors as early as the first day examined (day 6; Figs 2a and S3a and b). In the Donor 1 and Donor 3 derived organoids, notable E2-induced increases in *PGR* and *IHH* were observed on d9 compared to d6, while the level of *PGR* induced by E2 on d9 in the Donor 2 derived organoids decreased, and the level of *IHH* induced by E2 did not change relative to the levels observed on d6 (Figure 2a and Figure S3a and b). The transcripts were all calculated relative to levels observed in donor 1, revealing that, in general, estrogen induced all three transcripts more robustly in donor 1 derived organoids than in the other two. The expression of *ESR1* transcript was similar in all 3 sets of organoids (Figure S3c) but was lower in donor 2. In donor 2 samples, on day 9, induction of *PGR, IHH* or *GREB1* with E2 could be inhibited using the ESR1 antagonist, ICI 182,780, indicating an ESR mediated response (Figure 2b). Based on the responses observed, for comprehensive analysis of organoid transcriptomes, and ESR1 cistrome, we focused on samples from two donor derived organoids (donors 1 and 2) on day 9.

**Figure 2.**
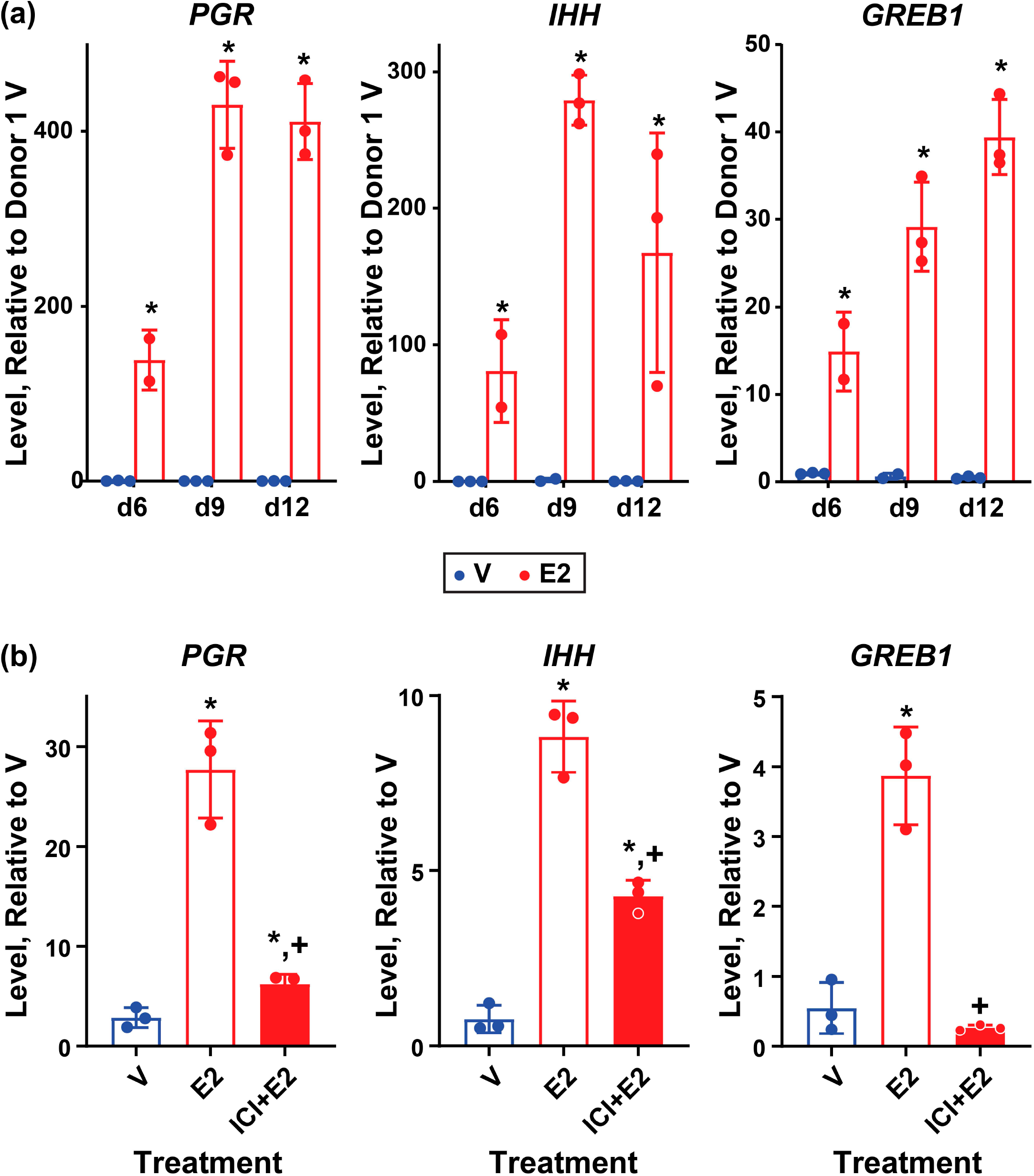
Estrogen responsive genes are induced in organoid cultures. a. RT-PCR of RNA isolated from donor 1 derived organoid cultures treated as described in methods and in Figure S2 with 0.1% ethanol vehicle (V) or with 10 nM estradiol (E2) 6, 9 or 12 days after initial plating (d6, d9, d12). Progesterone Receptor (*PGR*), Indian hedgehog (*IHH*) and Growth regulating estrogen receptor binding 1 (*GREB1*). Bar indicates mean, error bars indicate SD; * indicates p<0.05 vs. V based on 2-way ANOVA with Fisher’s LSD multiple comparisons test. N=3 for all but d6 E2 (n=2). b. RT-PCR of RNA isolated from donor 2 derived organoids 9 days after plating treated with 0.1% ethanol vehicle (V), 10 nM estradiol (E2) or with 1 µM ICI 182780+10 nM E2 (ICI+E2). Bars indicates mean, error bars indicate SD; * indicates p<0.05 vs. V; + indicates p<0.05 vs. E2 based on 2-way ANOVA with Fisher’s LSD multiple comparisons test. N=3 for all.

### Estrogen Regulates the Organoid Transcriptome

We used RNAseq to compare the transcriptomes of donor 1 and donor 2 derived organoid sets after treatment with vehicle or estrogen. Transcripts expressed in each donor were compared, indicating 10,652 genes were detected in both donors (Fig S4a). We then compared these genes to the 15,127 and 15,337 genes detected in proliferative and mid-secretory isolated endometrial epithelial cells (30), respectively (Fig. S4b), confirming that most of the organoid transcripts (76%) represent genes expressed in intact epithelium. Further analysis of the organoid transcripts revealed that, as was observed with the estrogen responsive genes evaluated by RT-PCR (Fig. 2 and Fig. S3), donor 1 derived organoids exhibited more robust estrogen responses than donor 2 ((Donor 1 E2/V 1907 DEG; Donor 2 E2/V 695 DEG; 2-fold, FDR < 0.05; Fig. 3a, Table S2a and b). The estrogen regulated organoid transcripts were analyzed using Ingenuity Pathway Analysis (IPA) revealing, as previously reported (47), that E2 treatment of organoids promotes formation of cellular protrusions/ciliagenesis (Table 2a and Table S2c). As in the whole endometrium, signals and regulators impacted overlapping processes, including chromatin modification, RTK mediated signals, and transcription (Tables 2b and c and Table S2c-e). Evaluation of the impact of gene regulations on signaling or regulatory pathways revealed activation of estrogen signaling, as well as Signal transducer and activator of transcription 3 (STAT3) and Transforming growth factor beta (TGFβ)- Smad family member (SMAD) mediated signals (Tables 2b and c), all of which have demonstrated importance in endometrial growth and function (48,49). The pattern of gene regulation of donor 2 is consistent with decreased activity of Wnt family member 5A (WNT5A), which is involved in polarity of epithelial cells during embryo implantation (50) (Table 2b). Activity of some functions or signals are selectively enriched in one donor, for example, Gamma-aminobutyric acid (GABA) and TP53 in donor 2, and AGT and SMAD in donor 1.

**Table 2.** Functions (a), Upstream Regulators (b) and Pathways (c) Enriched in Estrogen Regulated Organoid Genes

a. Biological functions. (overlap p value <0.05)
| **Function** | **Activation z-score** |  |
| --- | --- | --- |
|  | **Donor 1** | **Donor 2** |
| **Organization of cytoskeleton** | **4.50** | **4.88** |
| **Organization of cytoplasm** | **4.50** | **4.88** |
| **Microtubule dynamics** | **4.62** | **4.72** |
| **Formation of cellular protrusions** | **3.42** | **4.10** |
| **Formation of cilia** | **4.16** | **N/A** |
| **Transport of ion** | **3.44** | **N/A** |
| **Outgrowth of cells** | **-0.33** | **2.18** |

b. Upstream Regulators (overlap p value <0.05; activation z-score <-1.5 or >1.5)
| **process** | **Donor 1 regulators** |  | **Donor 2 regulators** |  |
| --- | --- | --- | --- | --- |
|  | **increased activity** | **decreased activity** | **increased activity** | **decreased activity** |
| **chromatin** |  | KDM1A |  | IKZF1 |
| **estrogen** | E2, ESR1 |  | E2 |  |
| **proliferation** |  |  | TP53 | mir-1 |
| **receptor signaling** | DEX, AGT, RXRA, PI3K |  | PI3K, DEX | GABA |
| **TGFβ signaling** | Smad2/3-Smad4 |  | Tgf beta |  |
| **cytokine signaling** | STAT3 |  | STAT3 |  |
| **transcription** | EHMT1, ERG | LDB1, LMO2 | ERG |  |
| **WNT signaling** |  |  |  | WNT5a |

c. Signaling Pathways (overlap p value <0.05; activation z-score >1.5)
| **process** | **Donor 1** | **Donor 2** |
| --- | --- | --- |
| **endometrial physiology** |  | **autophagy, relaxin** |
| **estrogen** |  | **ESR** |
| **receptor signaling** | **Adrenomedullin** | **Ephrin Receptor, Oxytocin, Androgen, Melatonin, Opioid, eNOS, VEGF-VEGFR** |
| **receptor tyrosine kinase signaling** |  | **G-β-γ, CRH** |

**Figure 3.**
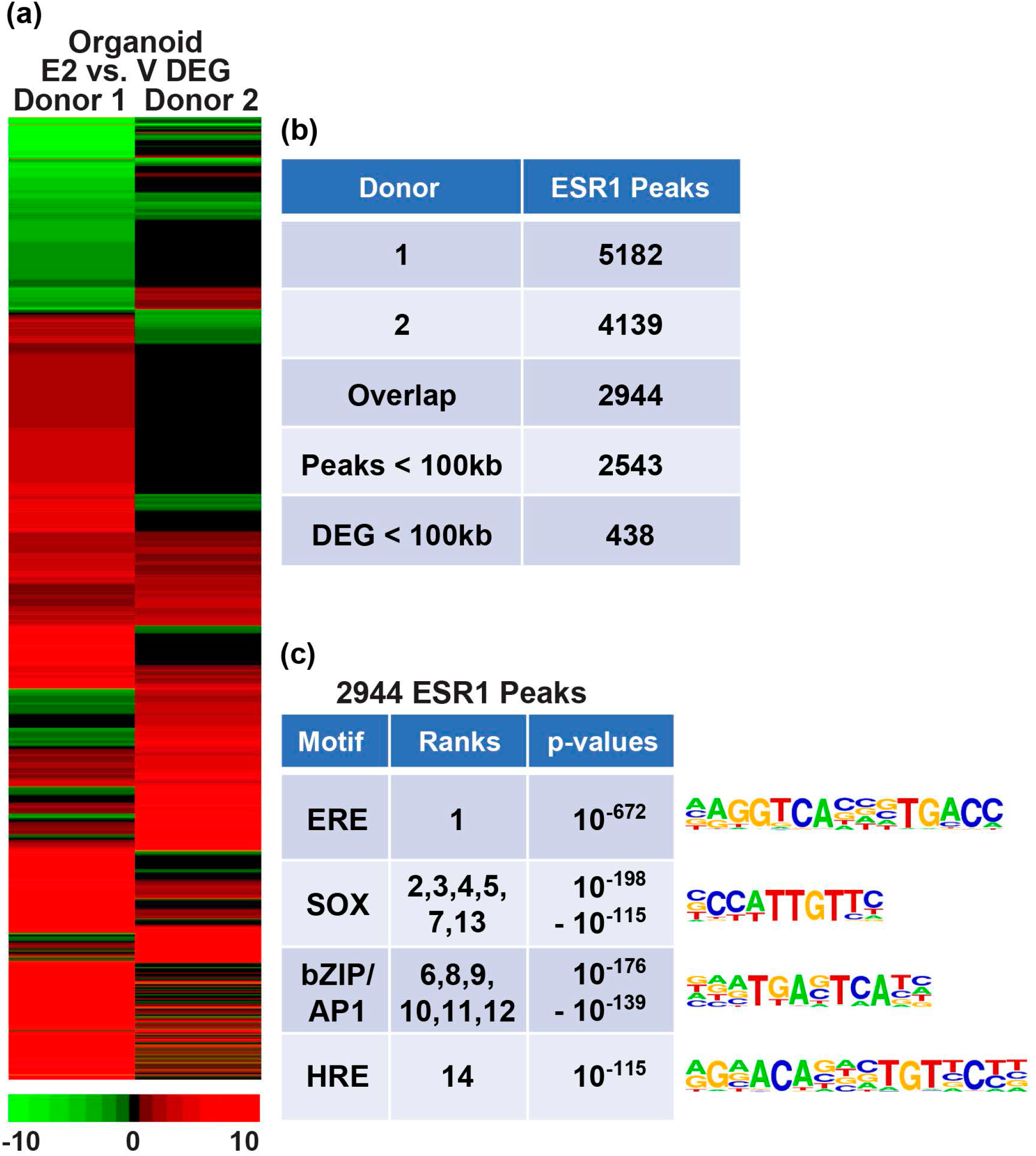
Human Epithelial Organoid ESR1 Transcriptome and Cistrome. a. Hierarchical cluster comparing E2 vs. V fold-changes of significantly regulated genes (2-fold, FDR p<0.05) in either donor. b. ESR1 ChIPseq of chromatin isolated from organoids. The number of peaks of each donor sample is indicated, as well as those shared by both donors. 2,543 of the 2,944 overlapping ESR1 peaks are within 100 kb of a gene. c. Summaries of the top 14 HOMER known motifs enriched in the 2,944 shared ESR1 peaks in organoids. The motif’s ranks, as determined by p-value, along with the range of p-values covered by the motif are indicated.

### Organoid ESR1 Cistrome

The gene regulation patterns revealed using RNAseq analysis indicated a robust response to E2. In order to better evaluate the direct effects of ESR1, sites of interaction with chromatin in organoids were evaluated by ESR1 ChIPseq. Comparable numbers of ESR1 peaks were identified in hormone treated organoids from either donor (Fig. 3b). For subsequent analysis, we focused on 2,944 ESR1 peaks shared by both donors. Examination of the locations of ESR1 peaks relative to transcripts shows that organoid ESR1 is more distal than the whole endometrial ESR1 (Fig. 1b), with a smaller proportion of binding in 5’UTR and exons and a larger proportion located in distal regions. As was observed in the proliferative endometrium, ERE was the most significantly enriched motif in the ESR1 peaks (Figure 3c). However, unlike in the whole endometrium, SOX motifs were significantly enriched. This likely reflects the role of the SOX transcription factor activity specific to epithelial cells. Like the whole endometrium, bZIP/AP1 motifs were enriched. The HRE motif, which interacts with PGR, AR and GR, was enriched as well. 2,543 of the 2,944 ESR1 peaks are within 100 kb of an annotated gene (Fig. 3b, Fig. S4c); 438 genes that are estrogen regulated in either donor organoid genes are < 100 kb from one of the 2543 ESR1 peaks (Fig. 3b; Table S3a). Pathway analysis of these 438 genes that are candidate ESR1 downstream targets indicates some of the signals and functions noted previously (Tables 3a and b and Table S3b and c), including formation of cellular protrusions, estrogen, STAT3 and TGFβ signaling, were enriched (Tables 3a and b). The pattern of gene regulation by these ESR1 targets revealed functions and regulators that were not apparent in the analysis of all the DEG (Table 2) including progesterone activation, and lipid concentration (Tables 3a and b).

**Table 3.** Functions (a) and Upstream Regulators (b) Enriched in Estrogen Regulated Organoid Genes That are Less Than 100 kb From ESR1 ChIPseq Peaks

a. Functions (overlap p value <0.05)
| **<u>Function</u>** | **Activation z-score** |  |
| --- | --- | --- |
|  | **<u>Donor 1</u>** | **<u>Donor 2</u>** |
| **Chemotaxis** | **2.34** | **0.6** |
| **Formation of cellular protrusions** | **2.26** | **1.28** |
| **Invasion of cells** | **1.75** | **1.46** |
| **Quantity of Ca<sup>2+</sup>** | **1.37** | **1.57** |
| **Concentration of lipid** | **-1.92** | **-1.22** |

b. Upstream regulator (overlap p value <0.05; activation z-scores <-1.7 or >1.3)
| **<u>process</u>** | **<u>Donor 1 regulators</u>** |  | **<u>Donor 2 regulators</u>** |  |
| --- | --- | --- | --- | --- |
|  | **<u>increased activity</u>** | **<u>decreased activity</u>** | **<u>increased activity</u>** | **<u>decreased activity</u>** |
| **chromatin** |  | let-7 |  | let-7 |
| **cytokine signaling** | STAT3, IL6, | IL22 | STAT3, IL6, | IL22 |
| **estrogen** | E2, ESR1 |  | E2, ESR1, |  |
| **growth factor/receptor tyrosine kinase signaling** | ERBB2 |  |  |  |
| **lipid** | ACOX1 |  | ACOX1 |  |
| **NR signaling** | NCOA2 |  |  |  |
| **progesterone** | P4 |  | P4 |  |
| **TGFβ signaling** | SMAD4 |  | SMAD4 |  |
| **transcription** | PPARG,<br>BCL6 |  | EOMES |  |
| **WNT signaling** | TCFL7, LGR4 |  |  |  |

To compare the impact of estrogen signaling in organoids and endometrium, ESR1 peak signal was compared in heat maps centered on organoid ESR1 peaks (Figure 4). Locations of organoid ESR1 are more like proliferative than to mid-secretory ESR1 peaks (Figure 4). To evaluate the impact of estrogen in whole endometrium and organoids, we compared the signaling and functions regulated by genes within 100 kb of ESR1 peaks in each. As with separate analyses (Tables 1 and 3) estrogen impacts multiple processes. Estrogen-E2 signaling is activated in both (Table 4b), whereas progesterone signaling is inactivated in proliferative endometrium and activated in organoids (Table 4b and Table S4). Some processes/functions are selectively enriched in organoids, such as formation of cellular protrusions, associated with cilia formation (Table 4a). Some are shared, such as ERBB2 activation, activation of lipid metabolic enzyme Acyl-CoA oxidase 1 (ACOX1), Dopamine, and AGT signaling, whereas others, including Forkhead box O1 (FOXO1) and Peroxisome proliferator activated receptor gamma (PPARG) signaling, are inhibited in the whole endometrium and activated in the organoids (Table 4b). Altogether this suggests that some processes intrinsic to the epithelial cells may be masked by whole endometrial signals and that culturing organoids reveals estrogen signaling processes that are not otherwise observable. Alternatively, the differences could reflect alteration of epithelial cell signals by their isolation and culture separate from the other cell types and other factors found in whole endometrium.

**Table 4.**
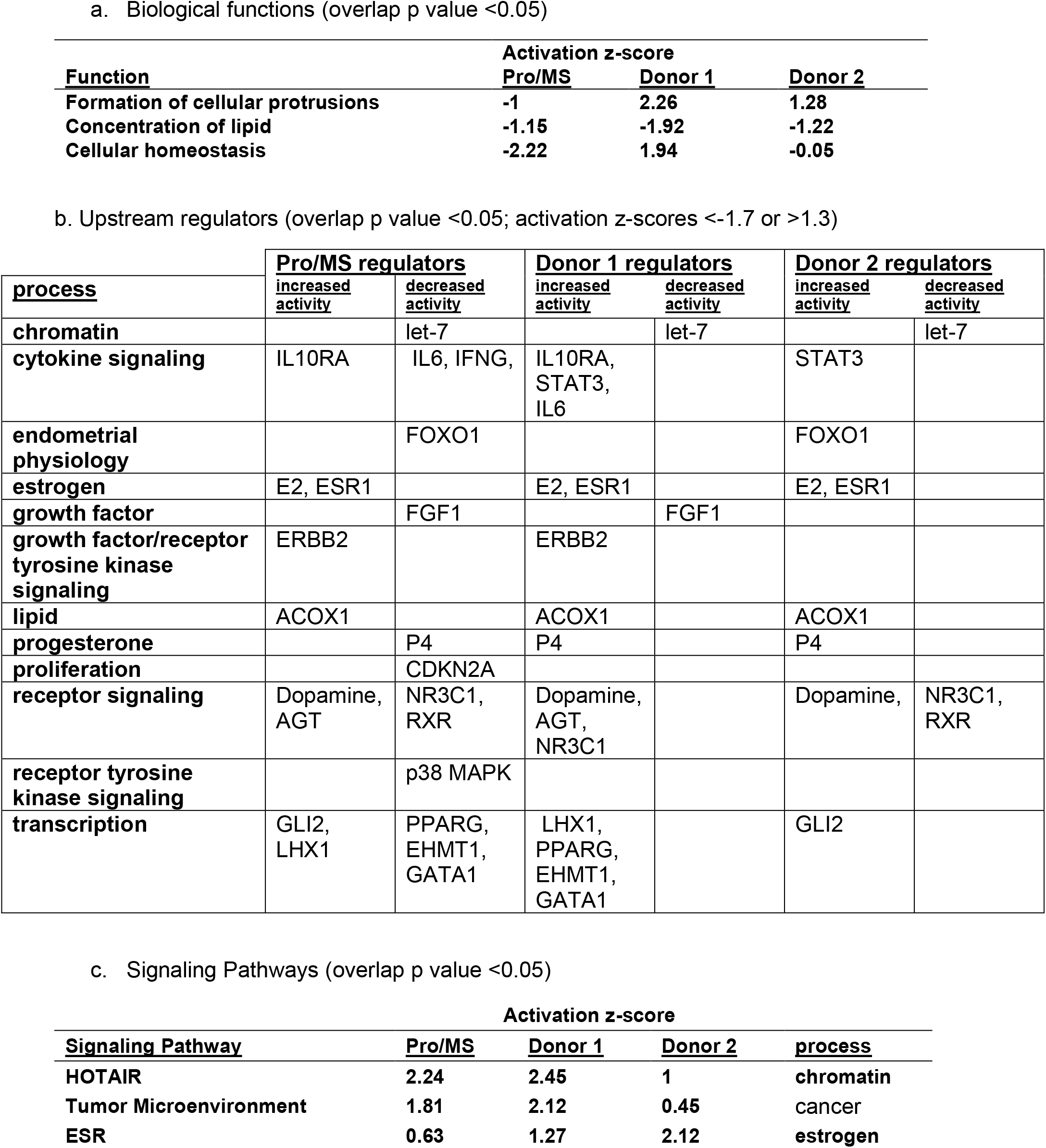
Functions (a), Upstream Regulators (b) and Pathways (c) Enriched in Proliferative vs Mid-secretory Endometrial and Estrogen Regulated Organoid Genes That are Less Than100 kb From ESR1 ChIPseq Peaks

**Figure 4.**
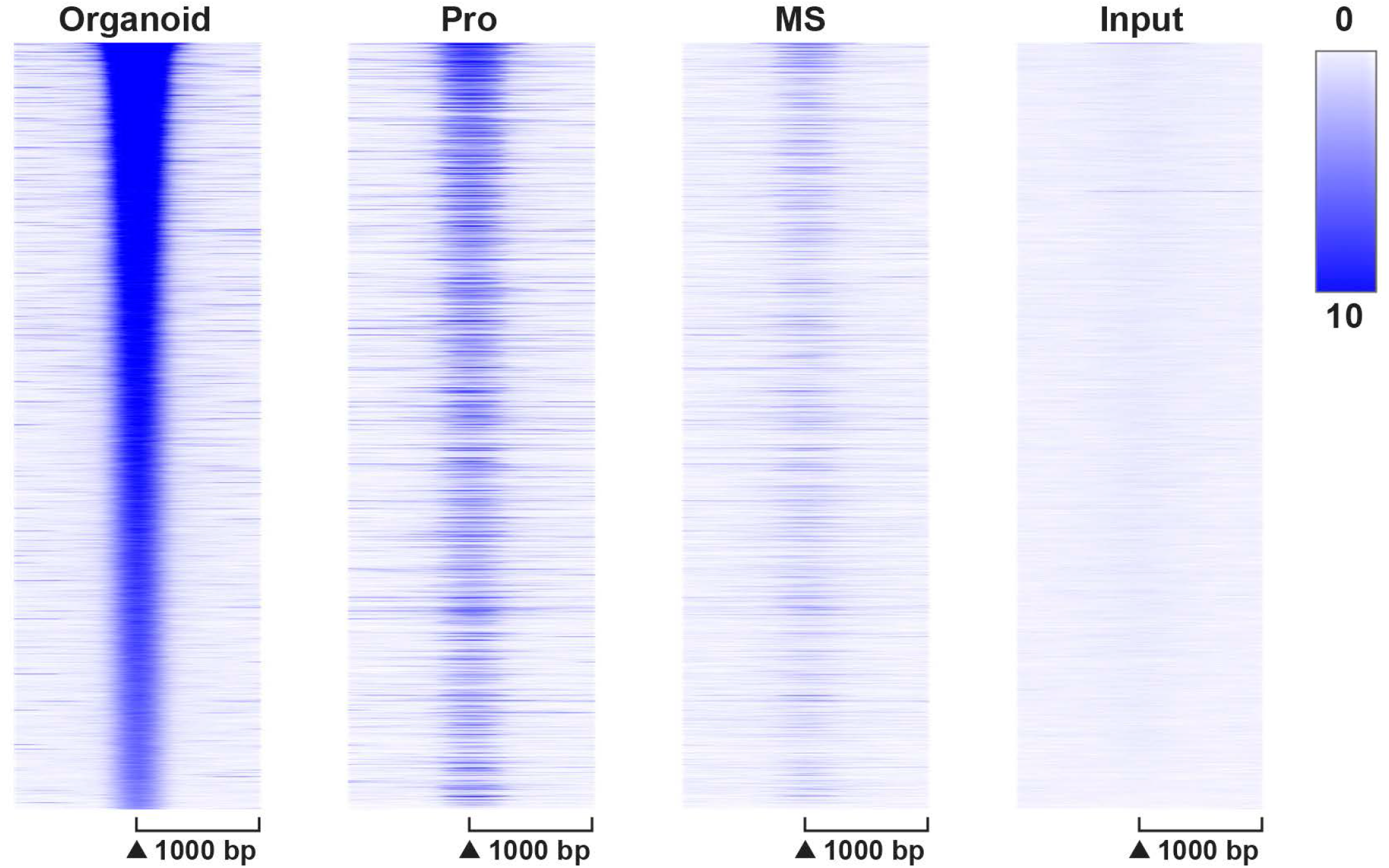
Organoid ESR1 Cistrome Resembles Proliferative Endometrium. ESR1 ChIPseq signal centered on locations of ESR1 peaks (+/-1000 bp) in organoid samples. Pro=proliferative whole endometrium; MS=mid-secretory whole endometrium.

Endometrial and organoid ESR1 peaks relative to *IHH and GREB1* genomic regions are shown in Fig. 5. Two ESR1 peaks are located 20 kb 5’ of the *IHH* TSS in organoids and in proliferative phase whole endometrium (Fig. 5a). Previous study described a similar region 19 kb 5’ of the mouse uterus *Ihh* gene (Fig. 5b), with demonstrated hormone dependent enhancer activity (51). ESR1 binds 70 kb 5’ of the *IHH* TSS in proliferative endometrium, and 100 kb 5’ of *IHH* 5’ in both proliferative and mid-secretory endometrium (Figure 5a). HiC from the ovariectomized mouse uterus (52) indicates that distal regions interact with the *Ihh* transcript (Fig. 5b). Multiple ESR1 peaks are present in a region beginning 50 kb 5’ of the *GREB1* gene (highlighted in yellow in Figures 5c) in organoids and endometrium. Several of the peaks in this region are more prominent in the proliferative sample as compared to the mid-secretory sample. The comparable region of mouse uterine *Greb1* was identified as a super enhancer with multiple ESR1 binding peaks in a previous study (52). In mouse uterus, HiC indicates interactions between the ESR1 binding super-enhancer region and the *Greb1* gene (Fig 5d). The pattern of ESR1 binding near these estrogen-regulated genes suggest potential enhancer regions involved in gene regulation by estrogen.

**Figure 5.**
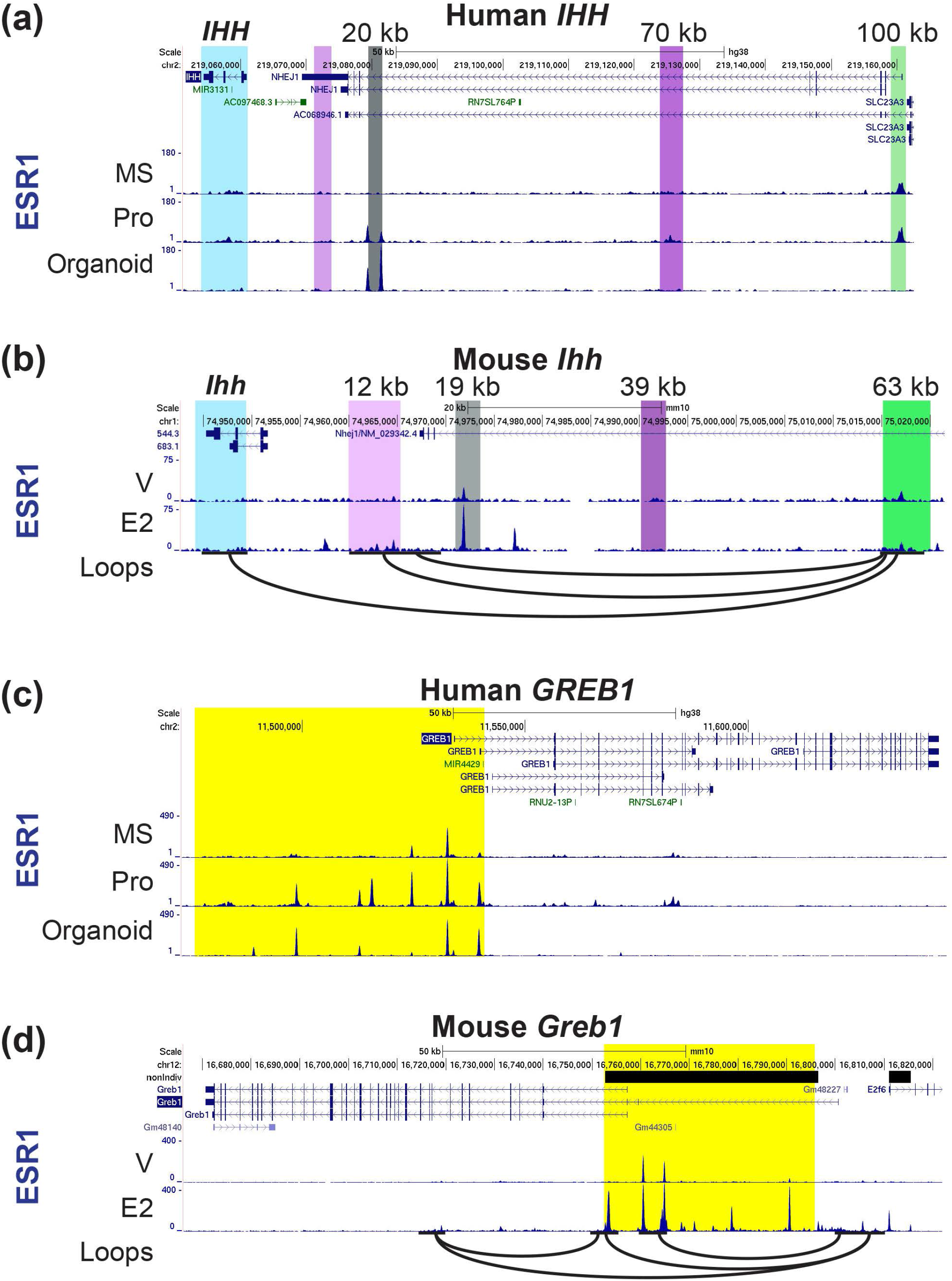
ESR1 peaks near human and mouse estrogen responsive transcripts *IHH*, and *GREB1*. For the human genes, ESR1 ChIP seq from mid secretory (MS) and proliferative (Pro) whole endometrium and from organoids is shown. For the mouse genes ESR1 ChIPseq from V or E2 treated whole uterus is shown. Interacting loops from mouse uterus HiC is also shown as black arcs. a. Human *IHH*. The transcript is highlighted in blue, regions with ESR1 peaks at 20kb 5’ of the *IHH* transcription start site (TSS) are highlighted in grey, at 70kb 5’ of *IHH* are highlighted in purple and at 100 kb 5’ of *IHH* are highlighted in green. b. Mouse *Ihh*. Regions comparable to the human gene are highlighted as described in a. c. Human *GREB1*. The ESR1-binding super-enhancer region is highlighted in yellow. d. Mouse *Greb1*. ESR1-binding super-enhancer is highlighted in yellow. Orientation on the genome browser view is the opposite of the human *GREB1*.

## Discussion

By combining cistromic and transcriptional data derived from endometrial biopsies or cultured epithelial organoids, we have shown details of estrogen mediated response in human uterine tissue. Our previous study described PGR ChIPseq analysis of endometrial biopsies and integrated it with RNAseq (30). Other published ESR1 ChIPseq studies utilized cultured endometrial stromal cells (53), uterine adenocarcinoma epithelial cell lines (23,54,55), endometrial tumors (24), or endometrial biopsies from infertile women (15). In these and other ESR1 ChIPseq analyses, most binding is observed either at distal enhancers or within genes, and motif analysis reveals that ERE is a highly enriched motif in ESR1 ChIPseq peaks from either stage ((18,19) Figs. 1b and c). One novel finding we report is preferential enrichment of homeobox factor (HOX) motifs in the mid-secretory ESR1 peaks (Fig. 1c). HOX proteins are transcription factors and chromatin modifiers involved in developmental patterning (56) that also facilitate stromal decidualization (57). Enrichment of HOX motifs was previously noted in mouse uterine ESR1 peaks that lacked ERE motifs (58). An ESR1 binding super-enhancer was described at the mouse *Hoxd* cluster (52), and is present in the proliferative human endometrium (not shown). Mouse models with disruption of *Hoxa 9,10,11 Hoxc 9*,10,11 and *Hoxd 9,10,11* exhibit impaired uterine development and function (59,60), and a mutation in HOXA11 was identified in a patient with a septate uterus (61). Together these observations reflect the importance of HOX factors to uterine biology.

bHLH motifs are enriched in proliferative and mid-secretory ESR1 peaks as well. HAND2 is a member of the bHLH family that modulates uterine stromal-epithelial signaling in mice (62). A recent study reported a single nucleotide polymorphism that interacts with *HAND2* as a potential causal factor for preterm birth (63). Enrichment of motifs that potentially bind the bHLH factor HAND2 within ESR1 peaks, together with its essential role in uterine biology, hint at an ability to impact overlapping genes and processes.

The “tethering” (indirect interaction) mode of ESR1 mediated transcriptional responses was derived from studies indicating ESR1 can interact with AP1 transcription factors to drive an AP1-luciferase reporter gene (64). Findings based on *in vitro* model gene systems did not account for chromatin architecture; major advances in our general understanding of cell specific gene regulation have highlighted the importance of modulation of chromatin accessibility, status of histone modifications, interaction between enhancers and promoters, and relative location within the nucleus (65). In this broader context, what appears as “tethering” in the simplified model likely indicates protein-protein interactions between ESR1 and other transcription factors (66) including bZIP (AP1), bHLH and HOX motif binding proteins, resulting in the enrichment of these motifs in the ESR1-binding sequences. Analysis of pathways impacted by estrogen in organoids or in proliferative endometrium included multiple chromatin modifiers (Tables 1, 2 and 3), pointing to the importance of chromatin dynamics in estrogen response. Transcripts for the factors that could bind DNA motifs enriched in endometrial ESR1 peaks (HOX, bZIP/AP1 and bHLH) are detected in the endometrial RNA seq data (Fig S5a), supporting their potential roles in estrogen signaling.

In organoids, ESR1 binding is more distal relative to genes than in the whole endometrium (Fig 1b). Whether this is a characteristic of epithelial cells or is a result of culturing isolated cells will be important to investigate. bZIP/AP1 motifs were enriched in organoid ESR1 ChIPseq peaks and organoid RNAseq datasets indicate expression of several AP1 family members (Fig S5b). Unlike the whole endometrium ESR1 peaks, SOX motifs were enriched in organoids. Multiple SOX transcripts are expressed, as reflected by the RNAseq (Fig S5b). A role for SOX in uterine epithelial cell function is supported by findings showing that SOX17 is detected in uterine epithelial cells in both mice and humans (51). In mice SOX17 binds to an enhancer 19 kb 5’ of *Ihh* that additionally binds multiple transcriptional regulators including ESR1 and PGR; this 19 kb enhancer is required for uterine *Ihh* expression (Fig 5b and (51)). bHLH motifs were highly enriched in proliferative and mid-secretory endometrium, but not in organoids (Figs 1c, 3c), consistent with the expression of HAND2 in uterine stromal cells (67). HOX motifs were not enriched in organoid ESR1 peaks and were the most significantly enriched motifs in mid-secretory endometrium, also consistent with the elevated expression of HOXA10 in mid-secretory stromal cells (68,69).

The organoid ESR1 cistrome is more like proliferative than the mid-secretory endometrium ESR1 cistrome (Fig. 4), and while estrogen stimulated transcriptional profiles of organoids compared to proliferative endometrium indicate some signaling processes are conserved, others differ. One limitation of the analysis is that the whole endometrium data relies on biological samples from women at proliferative and mid-secretory phases (estrogen dominant or progesterone dominant), which is not directly comparable to organoids treated with vehicle or estrogen. Genes that differ between the endometrial phases include ESR1 targets, but will also be influenced by progesterone, which is inhibiting estrogen activity in the mid-secretory phase. Enrichment of estrogen signaling activation in both the endometrial analysis and in organoids (Table 4b and c) indicates that the approach does capture estrogen targets in both systems and supports use of the organoid model to reflect endometrial epithelial cell estrogen response.

In both whole endometrium and in organoids, candidate ESR1 target signaling impacts chromatin remodeling associated processes (Tables 1-4), including HOTAIR, a long non-coding RNA encoded in the non-coding strand of the HOXC cluster. HOTAIR mediates epigenetic changes that repress genes by shuttling PRC2 and LSD1 to their targets (70). In multiple cancers HOTAIR promotes tumor growth, metastasis, migration and epithelial mesenchymal transition (70-72). HOTAIR is increased by estrogen in breast cancer cells (73), is associated with poor prognosis in endometrial cancer (74) and mediates endometrial cancer cell proliferation and invasion (75). Further investigation of estrogen regulation of this pathway in normal endometrium will be the focus of future work.

FOX01 signaling has been studied in mouse models, demonstrating expression in uterine epithelial cells and importance in uterine function (51,76,77), and show differences in enrichment between the epithelial cell organoids and the whole endometrium (Table 4b), highlighting the ability of the organoid model to reveal epithelial processes that are masked by the responses occurring in other endometrial cell types of whole endometrial samples. Our analysis also indicates estrogen impacts lipid and fatty acid metabolism, as ACOX1, a fatty acid oxidase, is activated and concentration of lipid is inhibited (Tables 4a,b). Lipid levels are vital for uterine receptivity and fetal development, and as sources for steroid hormone synthesis (78). Hormones impact lipid metabolism in mouse uterine epithelial cells (79), however the mechanisms and biological significance of estrogen/ESR1 regulation of endometrial lipid has not yet been well characterized.

Overall, our study provides novel details of ESR1 chromatin interactions underlying estrogen responses in the endometrium that can be further studied using epithelial cells cultured in organoids. We detect processes in the organoids that mirror those of proliferative endometrium, supporting their usefulness as a model to study estrogen responses intrinsic to epithelial cells.

## Acknowledgements

We are grateful to Jana Philips for coordinating endometrial sample donations for the studies.

## Data Availability

Original data generated and analyzed during this study are included in this published article or from GEO, as detailed in Results and Methods sections.

## Notes

This work was supported by the Intramural Research Program of the National Institute of Health: Project Z1AES103311-01 (F.J.D.), and Z1AHD008926 (AD), as well as NIH P01 HD106485 (S.L.Y.) and 1R01HD096266-04 (T.E.S.). The authors have no disclosures

### Competing Interest Statement

The authors have declared no competing interest.

