## Supplementary material for "The estrogen receptor α cistrome in human endometrium and epithelial organoids": Figs S1-S5 and legends

Figure S1

a. Summary of characteristics of ESR1 peaks and proliferative vs. mid-secretory DEG
b. Schematic illustrating how proliferative candidate estrogen target genes were derived.

Figure S2 Study design of organoid culture and estrogen response study.

a. Organoids were plated in Matrigel, allowed to form for 4 days, and then treated for 2, 5, or 8 more days with 0.1% ethanol vehicle (V) or with 10 nM estradiol (E2) added to media. For RNA isolation, the organoids were collected 6 hours after hormone treatments; for ChIP they were collected 1 hour after treatment.
b. Phase images of orgnoids.

Figure S3 Donor dependent organoid estrogen response

a. RT-PCR of RNA isolated from donor 2 and
b. donor 3 derived organoid cultures plotted relative to donor 1 (see Fig. 2a). Bars indicate mean, error bars indicate SD; * indicates $p<0.05$ vs. V based on 2-way ANOVA with Fisher's LSD multiple comparisons test. N=3 for all.
a. RT-PCR of *ESR1* comparing RNA isolated from V or E2 treated orgnoids derived from all 3 donors 9 days after plating. * indicates $p<0.05$ vs. V; # $p<0.05$ vs donor 1 based on 2-way ANOVA with Fisher's LSD multiple comparisons test. N=3 for all.

Figure S4 RNAseq of Organoids and Endometrial Epithelial Cells

a. Venn diagram of genes expressed (RPKM>1% of mean RPKM for all genes) in any sample (V or E2 treated).
b. Venn diagram of genes expressed in organoids and in epithelial cells isolated from proliferative or mid-secretory endometrium.
c. Schematic showing how candidate ESR1 regulated transcripts were derived.

Figure S5

a. FPKM values of bZIP/AP1, bHLH or HOX factors in proliferative (Pro) or mid-secretory (MS) endometrial samples.
b. E2/V ratios of f bZIP/AP1 or SOX factors in donor 1 or donor 2 derived organoids. “X” indicates the transcript was not detectable.

(a)

| | Proliferative | Mid-secretory |
|---|---|---|
| **ESR1 peaks** | **35156** | **8688** |
| **ESR1 peaks<100 kb from gene** | **31814** | **7931** |
| **Pro vs MS DEG** | **1628** | **1628** |
| **Pro vs MS DEGs <100 kb from ESR1 peak** | **902** | **394** |

(b)

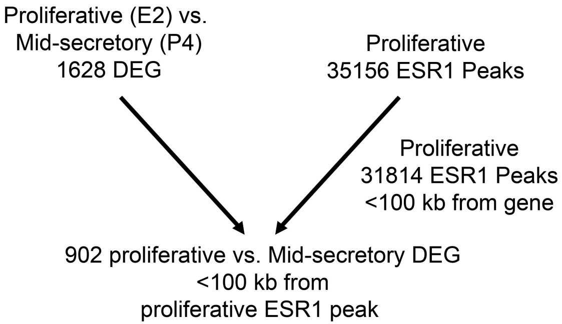

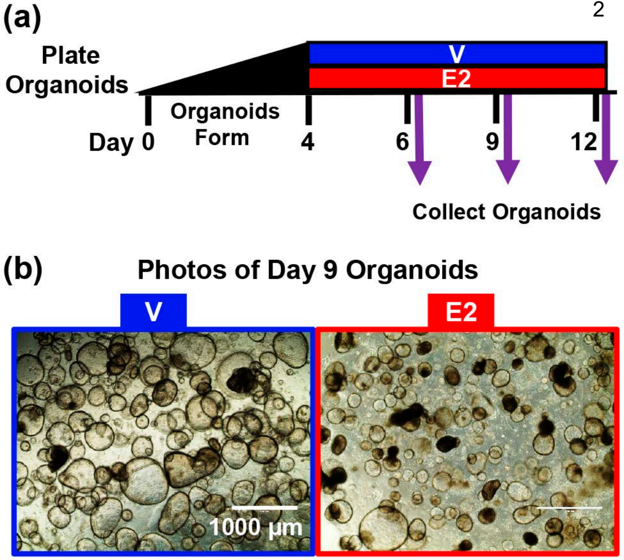
(a)
Plate Organoids
V
E2
Organoids Form
Day 0
4
6
9
12
Collect Organoids
(b)
Photos of Day 9 Organoids
V
E2
1000 µm

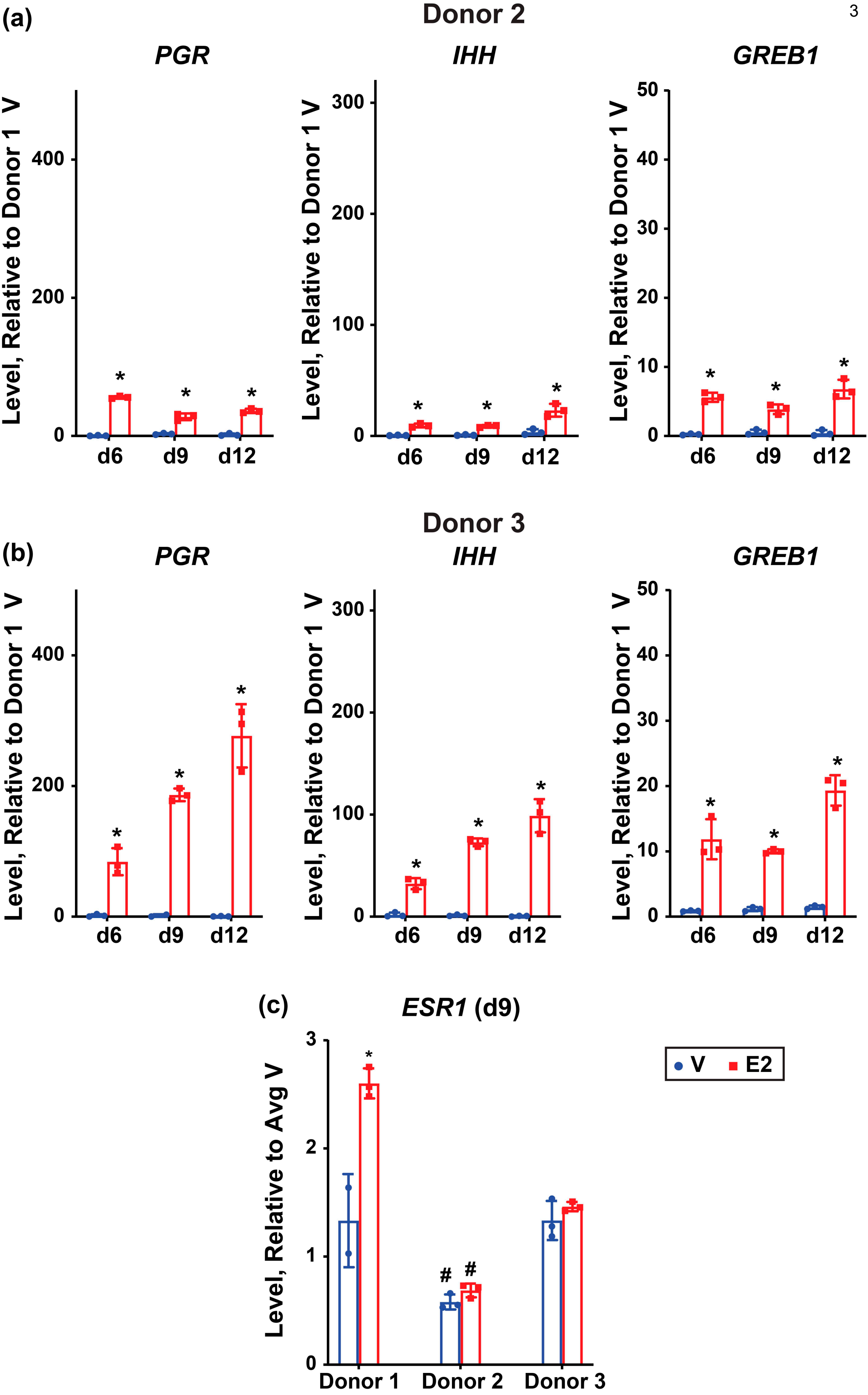
(a)
Donor 2
PGR
IHH
GREB1
Level, Relative to Donor 1 V
0
200
400
100
300
10
20
30
40
50
d6
d9
d12
*
(b)
Donor 3
PGR
IHH
GREB1
Level, Relative to Donor 1 V
(c)
ESR1 (d9)
Level, Relative to Avg V
0
1
2
3
#
Donor 1
Donor 2
Donor 3
V
E2

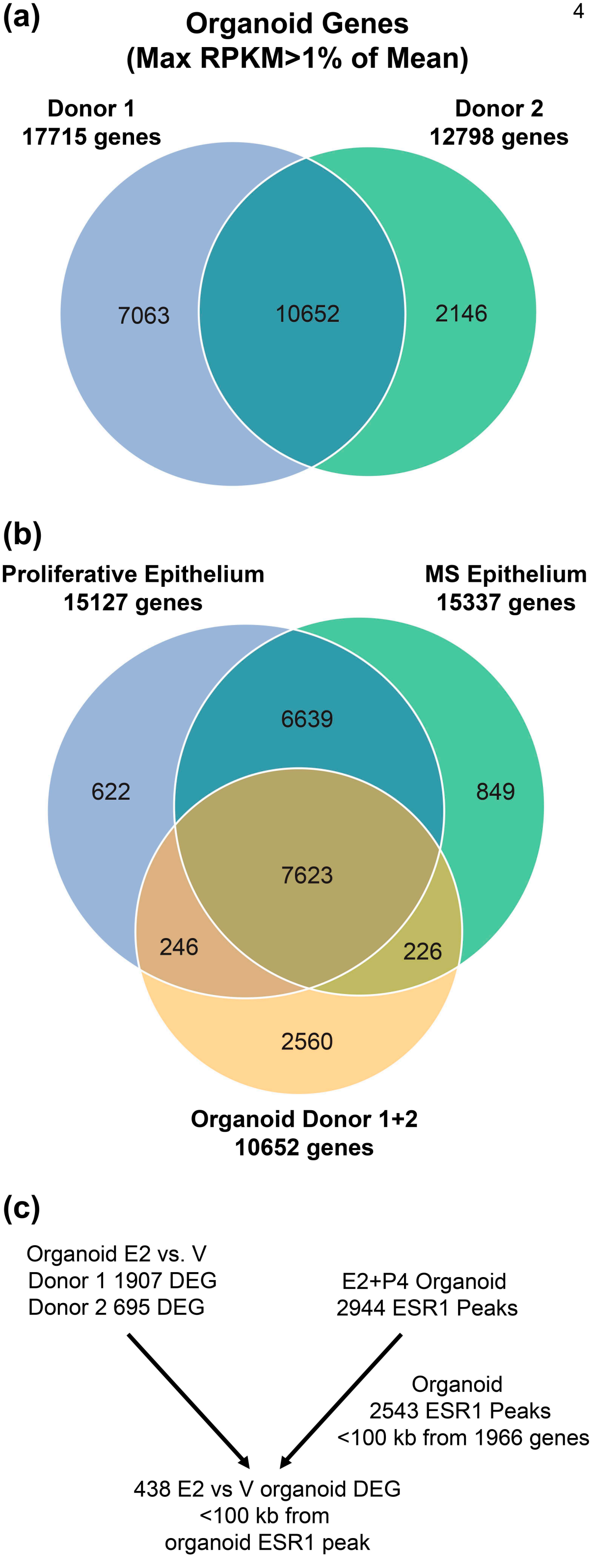
(a)
Organoid Genes
(Max RPKM>1% of Mean)
Donor 1
17715 genes
Donor 2
12798 genes
7063
10652
2146
(b)
Proliferative Epithelium
15127 genes
MS Epithelium
15337 genes
6639
622
849
7623
246
226
2560
Organoid Donor 1+2
10652 genes
(c)
Organoid E2 vs. V
Donor 1 1907 DEG
Donor 2 695 DEG
E2+P4 Organoid
2944 ESR1 Peaks
Organoid
2543 ESR1 Peaks
<100 kb from 1966 genes
438 E2 vs V organoid DEG
<100 kb from
organoid ESR1 peak

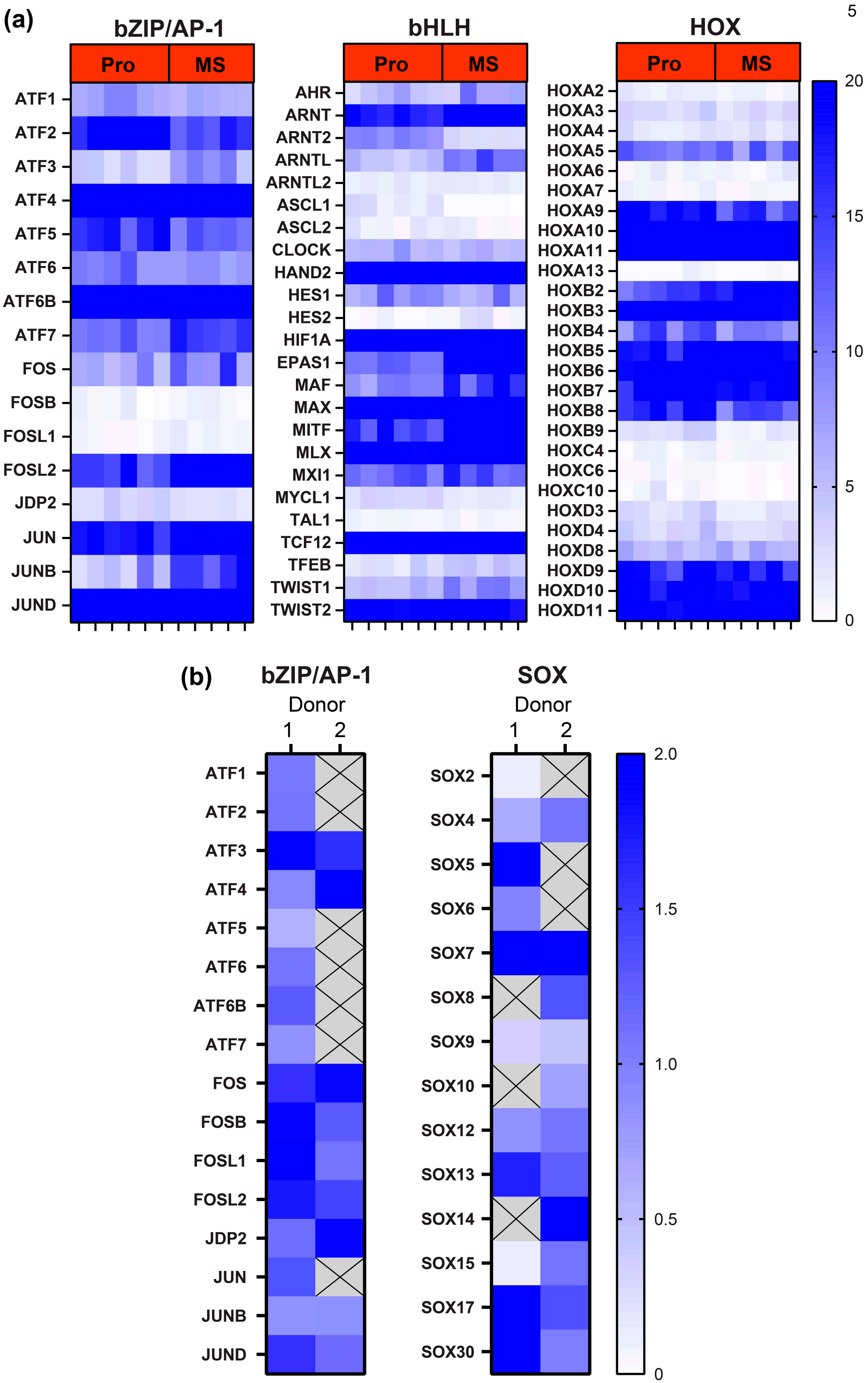
(a)
bZIP/AP-1
Pro
MS
ATF1
ATF2
ATF3
ATF4
ATF5
ATF6
ATF6B
ATF7
FOS
FOSB
FOSL1
FOSL2
JDP2
JUN
JUNB
JUND
bHLH
AHR
ARNT
ARNT2
ARNTL
ARNTL2
ASCL1
ASCL2
CLOCK
HAND2
HES1
HES2
HIF1A
EPAS1
MAF
MAX
MITF
MLX
MXI1
MYCL1
TAL1
TCF12
TFEB
TWIST1
TWIST2
HOX
HOXA2
HOXA3
HOXA4
HOXA5
HOXA6
HOXA7
HOXA9
HOXA10
HOXA11
HOXA13
HOXB2
HOXB3
HOXB4
HOXB5
HOXB6
HOXB7
HOXB8
HOXB9
HOXC4
HOXC6
HOXC10
HOXD3
HOXD4
HOXD8
HOXD9
HOXD10
HOXD11
5
20
15
10
5
0
(b)
bZIP/AP-1
Donor
1
2
SOX
SOX2
SOX4
SOX5
SOX6
SOX7
SOX8
SOX9
SOX10
SOX12
SOX13
SOX14
SOX15
SOX17
SOX30
2.0
1.5
1.0
0.5
0
